# The meta-epigenomic structure of purified human stem cell populations is defined at *cis*-regulatory sequences

**DOI:** 10.1101/007591

**Authors:** N. Ari Wijetunga, Fabien Delahaye, Yong Mei Zhao, Aaron Golden, Jessica C. Mar, Francine H. Einstein, John M. Greally

**Author notes:** These individuals made equal contributions to the manuscript (N. Ari Wijetunga, Fabien Delahaye).

## Abstract

The mechanism and significance of epigenetic variability in the same cell type between healthy individuals are not clear. Here, we purify human CD34+ hematopoietic stem and progenitor cells (HSPCs) from different individuals and find that there is increased variability of DNA methylation at loci with properties of promoters and enhancers. The variability is especially enriched at candidate enhancers near genes transitioning between silent and expressed states, and encoding proteins with leukocyte differentiation properties. Our findings of increased variability at loci with intermediate DNA methylation values, at candidate “poised” enhancers, and at genes involved in HSPC lineage commitment suggest that CD34+ cell subtype heterogeneity between individuals is a major mechanism for the variability observed. Epigenomic studies performed on cell populations, even when purified, are testing collections of epigenomes, or meta-epigenomes. Our findings show that meta-epigenomic approaches to data analysis can provide insights into cell subpopulation structure.

---

Variation in epigenetic marks defines specific cell types in an organism^1, 2^. Epigenome-wide association studies (EWAS) examine epigenetic variability within the same cell type or tissue in different individuals to assess the role of the epigenome in those individuals with a specific disease or other phenotype^3, 4, 5, 6^. In addition to epigenomic variability studied among different cell types in an individual or that in the same cell type among phenotypically different individuals, epigenomic variability occurring in the same cell type among healthy individuals is also now being studied^7, 8,, 9, 10, 11, 12, 13^. The mechanism and functional consequences of this type of epigenetic variability remain unclear. Such variability has been found in plants^14, 15^ and has been described as “inter-individual” differential methylation^12^ occurring at “epipolymorphic” loci that characteristically have intermediate DNA methylation levels^16^. The potential for stochasticity to drive at least part of this epipolymorphism of DNA methylation has been proposed^16^, and finds support from studies of allelic exclusion in the central nervous system of mouse^17^, monoallelic expression in neural stem cells^18^, and studies of heritability of DNA methylation in cloned ovarian carcinoma cells^19^. However, the proportions of genes at which these stochastic events are implicated is low (1–2%)^17, 18^, indicating that other processes are likely to be involved.

Underlying genetic polymorphism has been demonstrated to be a contributor to DNA methylation variability^10, 12, 13, 20^. Such genetic effects are unlikely to be the only influence, as monozygotic human twins^12, 20^ and inbred mice^8, 21^ also manifest epigenetic variability that cannot be attributed to DNA sequence differences. Some studies have linked DNA methylation variation with transcriptional consequences at nearby genes^8, 10, 12, 13^. Some of the variability observed in a study of peripheral blood leukocytes has been explained in terms of cell subtype effects^9^, although that study’s quantification of neutrophil, lymphocyte and monocyte percentages lacked the finer resolution cell subtype discrimination demonstrated in a later study to have effects on DNA methylation^22^. It has been shown that clinically normal cervical epithelial samples from women who proceed to develop cervical neoplasia within 3 years have increased variability of DNA methylation^11^. While this specific example reflects an underlying pathological process, epigenetic variability has also been proposed to be stochastic in origin and to influence normal phenotypic variability^8^. Supplementation of methyl donors in the diet of isogenic mice has been observed to increase the variability of DNA methylation in liver samples, suggesting to the authors a mechanism for disease or evolutionary selection^21^. The epigenetic variability observed in human CD14+ monocytes has been found to remain over the course of years, despite the short lifespan of these cells, indicating that the variability is encoded in leukocyte stem or progenitor cells^12^. Here we focus on using DNA methylation assays to define the loci with epigenetic variability in CD34+ hematopoietic stem and progenitor cells (HSPCs) purified from neonatal cord blood. We used the results of chromatin immunoprecipitation (ChIP-seq) studies of the same cell type by the Roadmap Epigenomics program to annotate the CD34+ HSPC genome empirically, so that we could define where epigenetic variability occurs in these cells, gaining insights into why the variability is occurring in seemingly-identical cell types from different healthy individuals.

## RESULTS

### Identifying variably DNA methylated loci in CD34+ HSPCs

We used two sources of DNA methylation data, one from the Roadmap Epigenomics program, publicly available reduced representation bisulphite sequencing (RRBS)^23^ data on mobilized CD34+ HSPCs from 7 adults, and the second generated by our group, using CD34+ HSPCs isolated from cord blood from 29 phenotypically normal neonates assayed using the HELP-tagging assay^24^. Despite the differences in how each of these assays measures DNA methylation, both showed increased variability at loci with intermediate methylation values (Fig. 1), consistent with previous observations^16^.

**Fig. 1.**
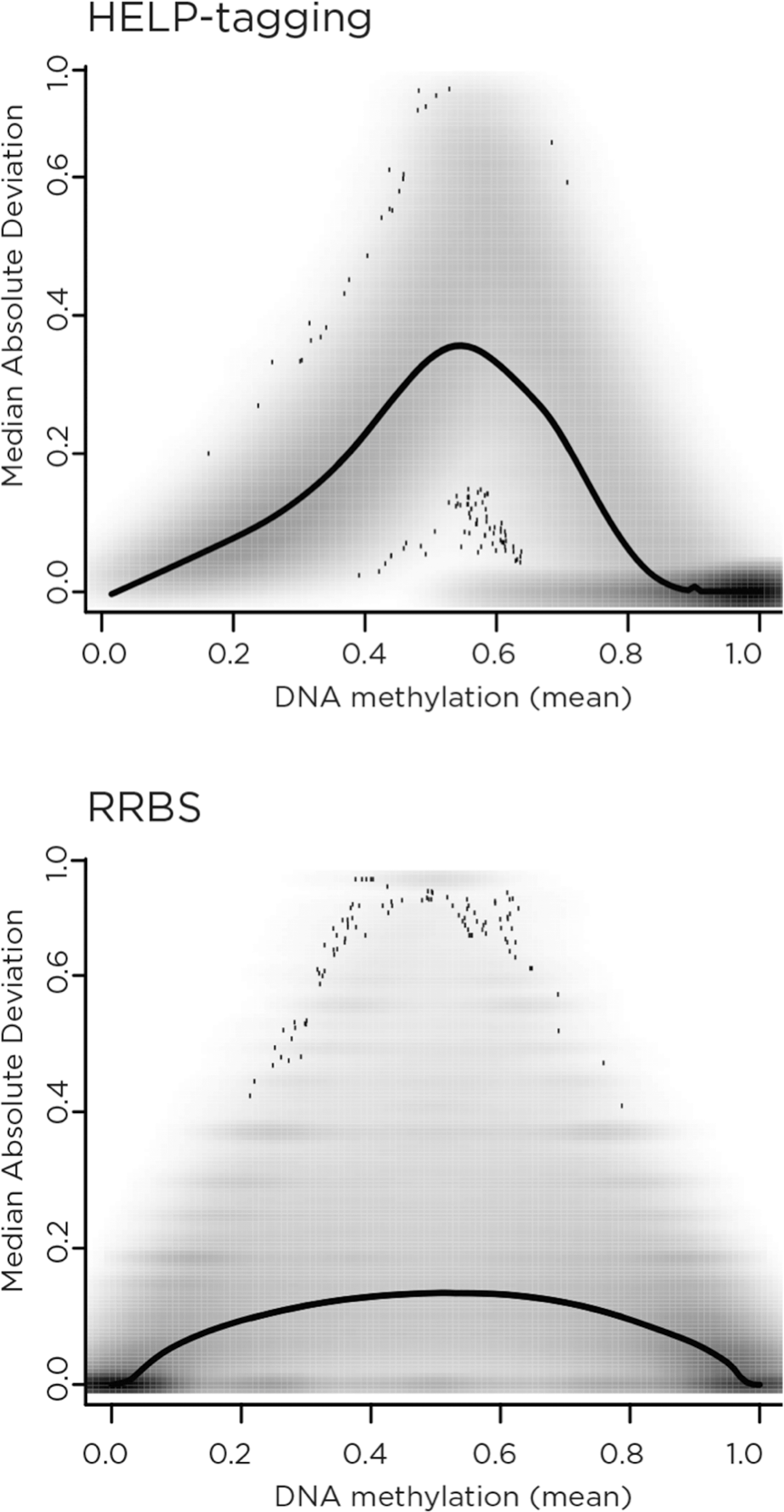
DNA methylation variability is increased at loci of intermediate methylation. The median absolute deviation (MAD) for DNA methylation values in CD34+ hematopoietic stem and progenitor cells (HSPCs) measured by HELP-tagging (top, 29 individuals) or reduced representation bisulphite sequencing (RRBS, bottom, 7 individuals) are shown as a function of mean DNA methylation across all of the samples tested. While HELP-tagging usually plots DNA methylation with a zero value to indicate complete methylation, we inverted the scale on this occasion to make the two plots comparable. The number of loci is reflected by the gray shading. The line shown indicates the mean MAD value, and reveals for both data sets increased variability of DNA methylation at loci with intermediate values.

We continued our analyses based on the HELP-tagging data, which are derived from a greater number of samples and from neonates, who have less potential for manifesting age-associated variability than adults^25^. As HELP-tagging is based on the use of methylation-sensitive restriction enzymes^24^, we were able to use the results from the methylation-insensitive MspI control enzyme to estimate the degree of technical variability, and a permutation analysis of the HpaII-derived data also showed enrichment of the observed variability over expected background levels (Supplementary Fig. 2a). A number of loci with differing degrees of variability were chosen for bisulphite PCR, using 7 of the samples that had been tested using HELP-tagging as well as 8 independent samples. These amplicons were combined for each individual and used to generate Illumina libraries, allowing targeted massively-parallel sequencing of the bisulphite-converted DNA. The results confirm that DNA methylation variability is enriched at loci with variability measures above the threshold attributable to technical variability or chance (Fig. 2b).

**Fig. 2.**
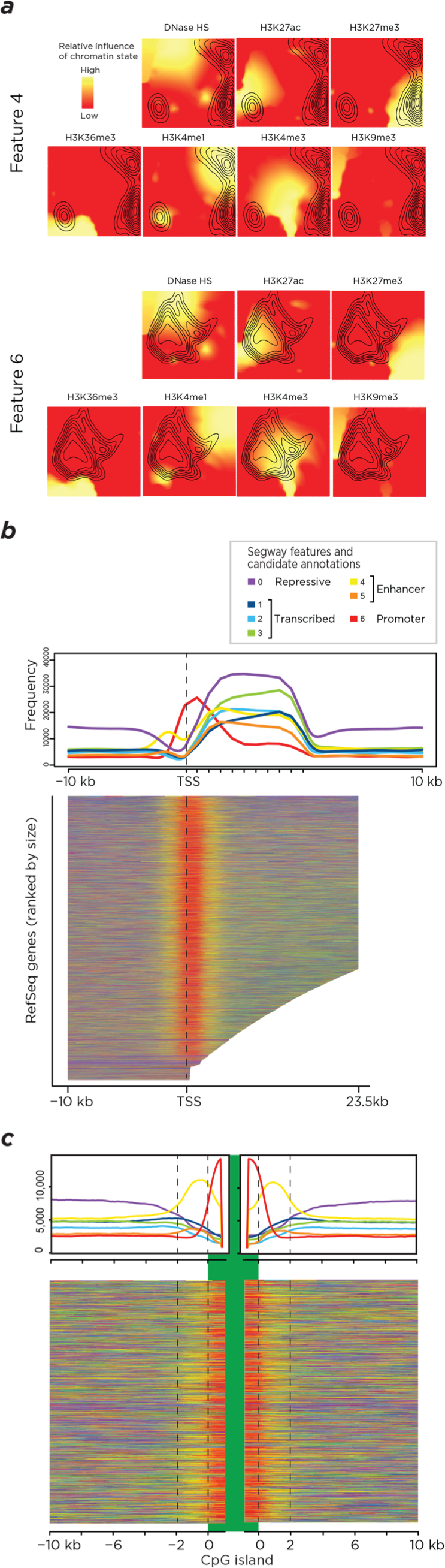
Empirical annotation of the CD34+ HSPC genome based on chromatin features reveals candidate *cis*-regulatory element locations. Panel (a) shows a contour plot of the regions within the self-organizing map (SOM) where Segway features 4 (above) and 6 (below) enrich, showing feature 4 to be composed of loci where H3K4me1 and H3K27me3 occur, while the loci composing feature 6 contain the H3K4me3 and H3K27ac modifications. Consistent with these findings, panel (b) shows feature 6 (red) to be enriched at the transcription start site for a metaplot (top) and a heat map (below) of all RefSeq genes, indicating promoter characteristics, while feature 4 (yellow) flanks this region and is consistent with enhancers in a poised state. In panel (c), similar metaplot (top) and heat map (below) representations of the 2 kb flanking CpG islands demonstrate strong enrichment in feature 4, indicating that these “CpG island shores” in fact represent candidate enhancers in this cell type.

### Inferring the effects of DNA sequence variability

To test whether the variability we observed could be accounted for by genomic sequence polymorphism, we segregated the variability at loci overlapping common single nucleotide polymorphisms (SNPs, minor allele frequencies ≥1%, 7.6% of sites tested) from the remaining majority of the genome. A Kolmogorov-Smirnov test (K-S test) showed significantly increased levels of variability of DNA methylation at these polymorphic loci (p<2×10^−16^), indicating that genetic influences are contributing to the variability observed (Supplementary Fig. 3). There are two ways that local sequence variability can influence DNA methylation variability. The site being tested in the DNA methylation assay can itself be a sequence variant, as cytosine to thymine transitions at CG dinucleotides represent a frequent source of SNPs due to the increased mutability of methylcytosine^26^, leading to the failure of methylation-sensitive restriction enzymes to cut or the misleading appearance of bisulphite-mediated conversion at these sites. The second mechanism is for SNPs *in cis* to the tested site influencing DNA methylation, as has previously been shown^27^. We find that the K-S test is no longer significant (p=0.1563) at sites tested even within the immediate flanking 10 bp of the common SNPs (Supplementary Fig. 3). We infer that while genetic variability is influential at the tested loci themselves, there exists a substantial amount of epigenetic variability in the remaining majority of loci in the genome, and that local genetic polymorphism is not likely to be the sole cause of the epigenetic variability observed, consistent with the conclusions of prior studies^8, 12, 20, 21^.

**Fig. 3.**
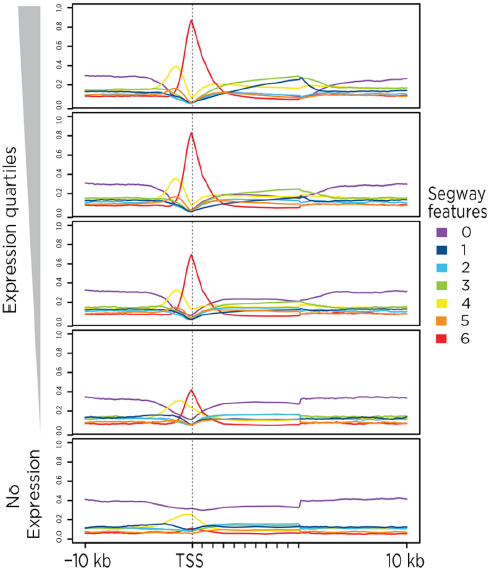
Transcriptional relationships of Segway features. A RefSeq metaplot for the Segway features divided by expression quantile shows that features 1–3 enrich in the bodies of genes as transcription increases, at the expense of feature 0, which appears to represent repressed chromatin. Feature 6 is strongly enriched at canonical transcription start sites, flanked by an enrichment of feature 4 and, to a lesser extent, feature 5, which have chromatin signatures indicative of enhancer function.

### Mapping functional elements in CD34+ HSPCs

To determine whether epigenetic variability was occurring at regulatory sites with possible functional consequences, we took advantage of public chromatin mapping data for CD34+ HSPCs generated by the Roadmap Epigenomics program. The DNase hypersensitivity and ChIP-seq data create combinatorial patterns that have previously been exploited to define functional elements in the genome^28^. We processed the Roadmap data using an adaptation of an imaging signal processing algorithm^29^ to define the locations of chromatin constituents with minimal data transformation (Supplementary Fig. 4). These chromatin constituent locations were then used to generate a self-organizing map (SOM)^30^, and to map candidate regulatory elements using the Segway algorithm^28^ (Supplementary Fig. 5). The individual Segway features were then overlaid as contour plots onto the SOM, which clusters in two dimensional space loci with similar genomic characteristics, allowing intuitive visualization of the major contributors to each feature (Fig. 2a and Supplementary Fig. 6). Of the multiple chromatin states for which each feature is enriched, feature 6 has the H3K4me3 enrichment indicating promoter function, features 4 and 5 both have marks indicative of enhancer function (H3K4me1 and H3K27ac, respectively), features 1–3 have the H3K36me3 enrichment typical of transcribed sequences while feature 0 in enriched for heterochromatic marks (H3K9me9 and H3K27me3). We also created a metaplot of these new annotations relative to all RefSeq genes in the genome, showing that Segway feature 6 is strikingly enriched at transcription start sites (TSSs), flanked by enrichment for feature 4 and, to a lesser degree, feature 5 (Fig. 2b). Features 1–3 are enriched in gene bodies and feature 0 at intergenic sequences. Statistical testing of the enrichment of features 4 and 6 in their windows of peak frequencies compared with their distributions over all RefSeq genes and flanking regions showed significance (p<0.001 for each). CpG islands and their immediate flanking sequences have previously been related to “stochastic” DNA methylation variability^8^ and gene expression regulation^31^. The Segway annotations demonstrate that while the bodies of CpG islands are enriched for the candidate promoter (feature 6) sequences, the ±2 kb flanking region, generally described as its “shore”, is strikingly enriched for feature 4 (Fig. 2c). Both achieve statistical significance (p<0.001) when compared with their distributions over all CpG islands (feature 6) or flanking regions (feature 4). Finally, stratifying the RefSeq genes by expression quartile in CD34+ HSPCs reveals the transcriptional dependencies of the Segway annotations (Fig. 3). We conclude that the Segway annotations define candidate promoters (feature 6), enhancers (features 4 and 5), transcribed regions (features 1–3) and repressed chromatin (feature 0) for CD34+ HSPCs.

**Fig. 4.**
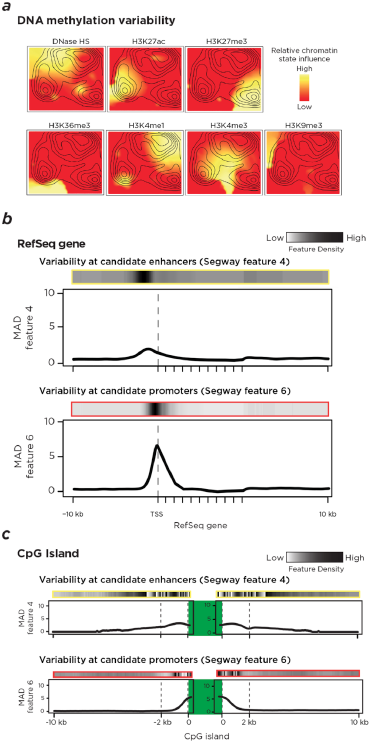
DNA methylation variability is enriched at candidate enhancers and promoters at transcription start sites of RefSeq genes and at CpG islands and shores. Panel (a) shows that enrichment of variability of DNA methylation is marked at loci with H3K4me1, H3K27ac and H3K27me3 marks in particular. Panel (b) shows a RefSeq metaplot with feature density indicated by the gray shading above and within the graph, and mean variability for features 4 (yellow, top) and 6 (red, bottom) depicted, with increased variability distributing where each mark is maximally located. The significance of the enrichment is shown at the depicted peak p value location. Analysis of CpG islands (c) shows variability in flanking regions (shores) associated with the presence of feature 4.

**Fig. 5.**
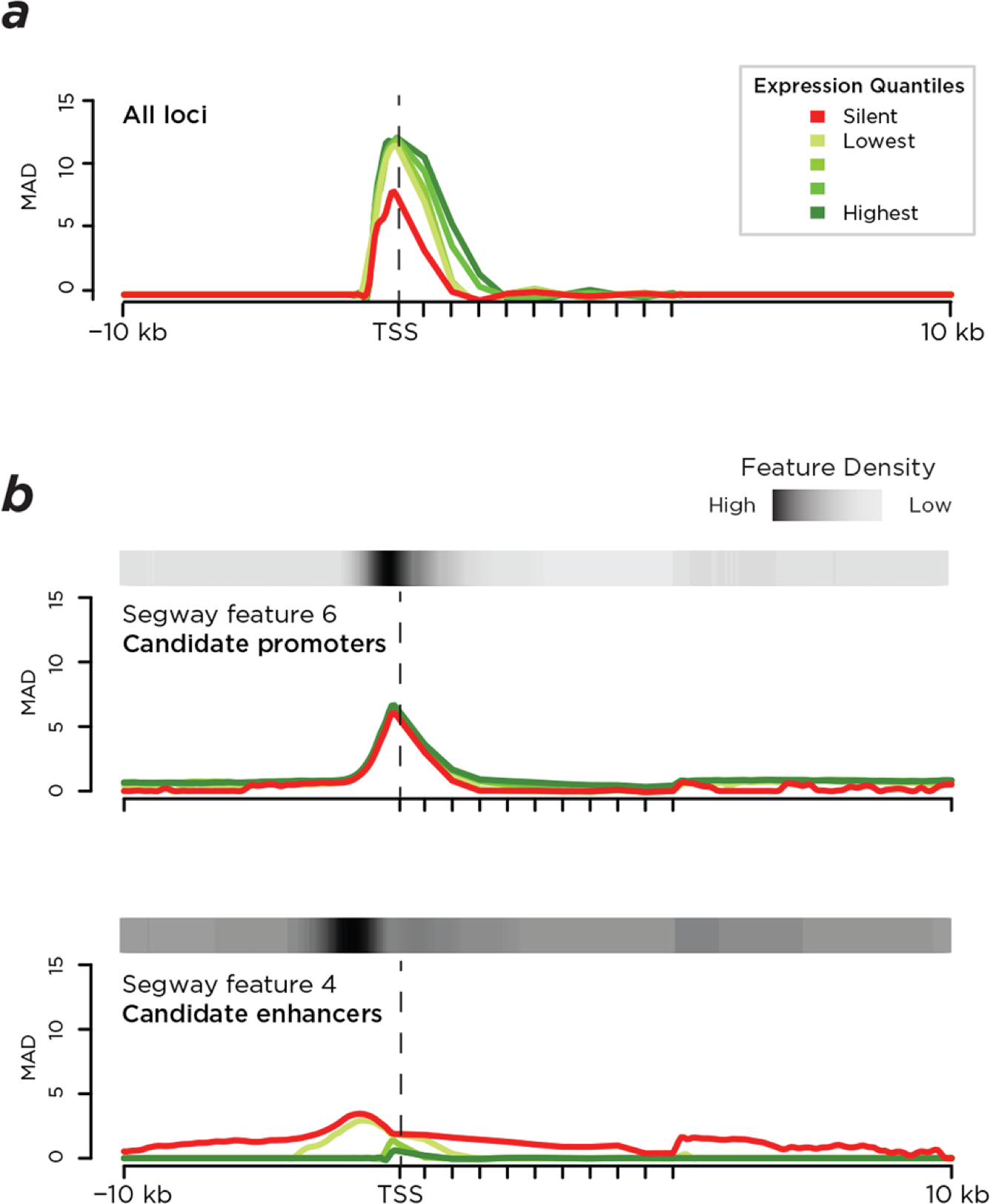
Variability of DNA methylation at candidate enhancer sequences discriminates genes expressed at lower levels. Panel (a) shows the overall pattern of DNA methylation variability at RefSeq genes broken down by expression quantile, showing differences at silent compared with expressed genes at transcription start sites. (b) No such differences are measurable when testing candidate promoters (Segway feature 6, top), whereas candidate enhancers (Segway feature 4, bottom) show increased variability for DNA methylation for genes that are either silent or expressed at the lowest quartile.

**Fig. 6.**
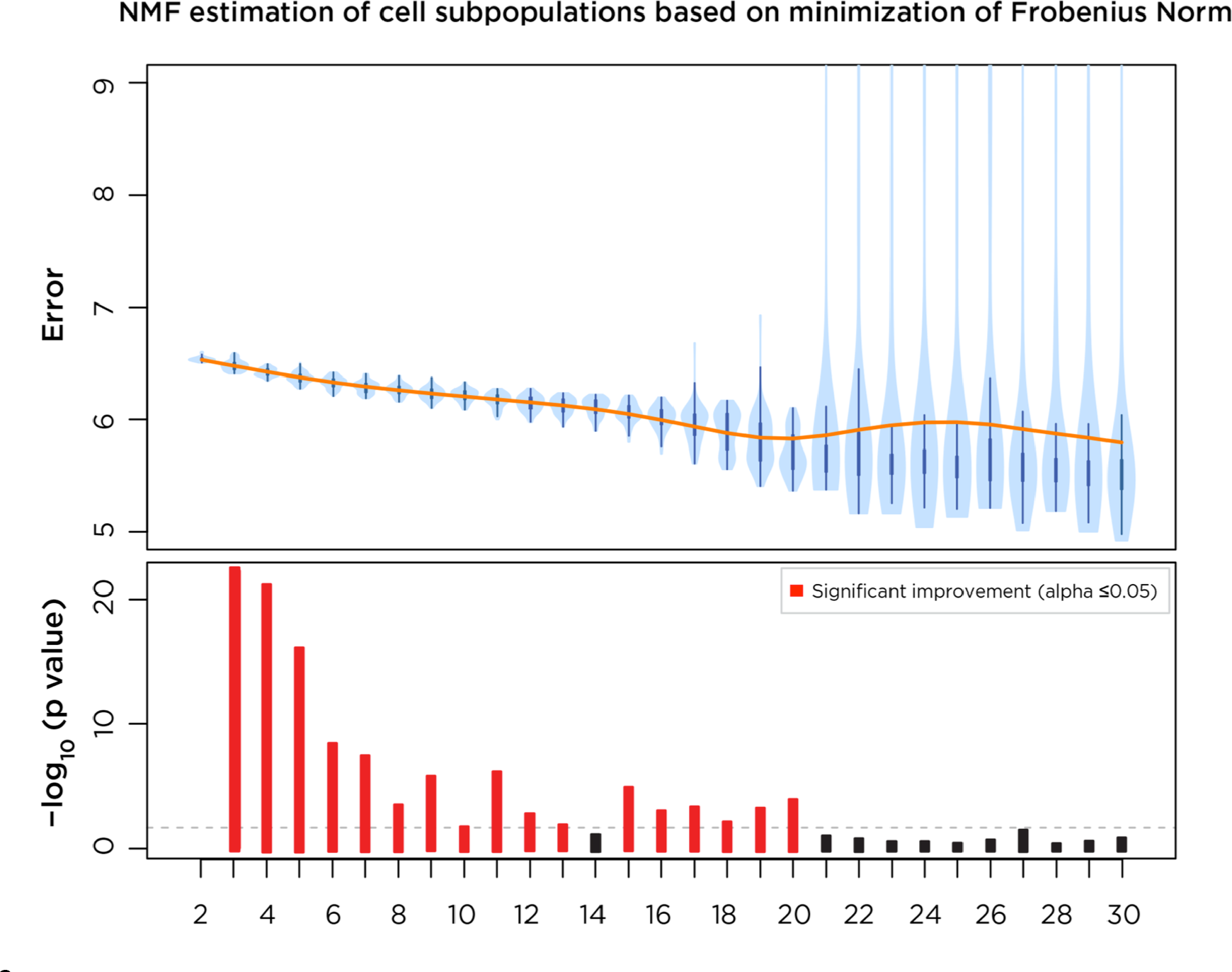
Non-negative matrix factorization (NMF) of DNA methylation profiles shows evidence for 13–20 subpopulations within the CD34+ HSPC population. The upper plot shows a smooth spline (orange) and value distribution (blue) for the Frobenius norm as a function of increasing cell subpopulation number, with the lower plot representing the p value testing (two sample t-test) whether the difference between the successive simulations is significant. We observe two points at which the subsequent change is insignificant, at values 13 and 20, suggesting that the number of subpopulations differing in DNA methylation profiles within the CD34+ HSPC population is within this range.

### Variable DNA methylation is enriched at functional elements

With the genome annotated for functional elements in a cell type-specific manner, we then tested the associations between genomic annotations and the loci with increased variability in DNA methylation. In Fig. 4a, we show the strongest associations for highly variable loci within clustered SOM space to be with H3K27ac, H3K27me3 and H3K4me1. Fig. 4b also shows enrichment of variability at the TSS of RefSeq genes for feature 6 and immediately upstream at feature 4, both significant at p<0.001. Fig. 4c shows enrichment in variability at the proximal part of CpG island shores for feature 6 and more extensively into the CpG island shore for feature 4, both also significant at p<0.001. A complementary SOM analysis using the published ChromHMM annotations of the human genome^32^ reveals consistent results (Supplementary Fig. 9). DNA methylation variability is therefore enhanced at candidate *cis*-regulatory sequences (promoters and enhancers) and the epigenetic variability previously observed for CpG island shores^8, 31^ is reflective of this general characteristic of enhancers. Common SNPs are not enriched in density in any of the features (Supplementary Fig. 10) and therefore are unlikely to be the major reason for selective enrichment of epigenetic variability in these specific genomic contexts. If the variability of DNA methylation occurs at loci with potential transcriptional regulatory properties, it raises the question of whether variability occurs selectively near genes with specific transcriptional activities. We find that all levels of expression have comparable levels of epigenetic variability at promoter sequences. Genes expressed at the lowest levels in the genome are those with selective enrichment of epigenetic variability at nearby candidate enhancers (Fig. 5), a significant inverse relationship (p=10^−8^) using the Jonckheere trend test.

### Variable DNA methylation is enriched at hematopoietic genes

The finding of increased variability at loci with intermediate values of DNA methylation (Fig. 1) cannot be explained by individual cells having intermediate methylation values, as DNA methylation is either present or absent on an allele, so a cell can only have 0, 50 or 100% methylation at a locus with two parental alleles in a diploid cell. For one individual to have an intermediate value such as 30% and another 60% methylation at the same locus, there has to be, within the pool of cells tested, differing subpopulations of methylated alleles present. The parsimonious explanation for such allelic subpopulation differences is that different proportions of cell subtypes are present in the individuals tested. The CD34+ HSPC is a well-studied cell type, recognized to have ∼15 cellular subtypes bearing the surface marker in different lineages and stages of early hematopoietic differentiation^33^. If these subtypes differ in proportion between the 29 subjects tested, the loci where DNA methylation patterns are associated with specific CD34+ cell subtypes would be expected to be the most variable, whereas genes expressed in all cell types (housekeeping genes) should be invariant in terms of DNA methylation. We show in Supplementary Table 6 and Supplementary Fig. 11 that DNA methylation variability at candidate enhancers is enriched for leukocyte-specific networks but not housekeeping genes, consistent with a CD34+ cell subtype model. Candidate promoters, which have comparable levels of DNA methylation variability at all expression levels (Fig. 5), have, as expected, equivalent levels of variability for housekeeping and hematopoietic genes.

### Using variability information to quantify cell subtypes

Variability of gene transcription levels in cell samples from multiple individuals has allowed patterns to be identified that predict the numbers of cell subtypes present. We adapted one of the approaches used for these transcriptional variability studies, Nonnegative Matrix Factorization (NMF)^34, 35^, to our DNA methylation variability data to estimate the number of cell subtypes in our purified CD34+ HSPCs. In Fig. 6 we show the NMF output to predict ∼13–20 cell subtypes, consistent with the ∼15 distinct types of cells that have previously been described to express the CD34 cell surface marker^33^.

## DISCUSSION

This study explored the epigenomic variability between normal healthy individuals occurring in a purified cell type, homogenous for the same cell surface marker. In common with prior studies, we found DNA methylation to vary between individuals, especially for those loci with intermediate DNA methylation values^16^, and increased variability at regulatory regions near the genes expressed at the lowest levels in the genome^12^. Our empirically-based annotation of regulatory elements in the CD34+ HSPC genome allowed us to define candidate promoter and enhancer sites, revealing the latter in particular to have DNA methylation variability associated with the lowest levels of gene expression. The genes at which this enhancer-associated variability is enriched are those encoding proteins with properties associated with leukocyte function. Because of the necessity for intermediate DNA methylation values to require allelic subpopulations with distinct methylation states, we interpret the epigenomic variability to be due to DNA methylation-sensitive enhancers changing their epigenetic states as nearby genes switch their transcription on or off in different CD34+ HSPC cell subtypes. The presence of differing proportions of these cell subtypes in the different individuals studied, and the consequent differences in allelic proportions of methylated DNA at these *cis*-regulatory sites, together drive the variability observed.

In keeping with this model of allelic and cell subtype heterogeneity, the chromatin signature of feature 4, that of “poised” enhancers^36, 37^, is defined by the co-occurrence in the genome of activating H3K4me1 and repressive H3K27me3 marks. As cells commit to the use of the enhancer, the locus is activated and marked by H3K27ac accumulation^38^ and the loss of H3K27me3^36^. Implicit in the idea of a poised regulatory element is its capability to commit to an active or inactive state by choosing one of two pre-existing states encoded in the nucleosome, as demonstrated by sequential ChIP experiments defining bivalent chromatin domains in pluripotent mouse cells^39^. Poised enhancers have not been reported to have been tested by sequential ChIP, making it possible that the activating and repressive marks are not encoded within the same nucleosome, but occur on different alleles in the cell population studied. This would be consistent with the presence of distinct subtypes of cells within the population tested by ChIP-seq when defining these poised enhancers. Our finding of variability and intermediate levels of DNA methylation at these loci with the features of poised enhancers is suggestive of epigenetic heterogeneity among the alleles in the cell population. The presence of the activating and repressive marks of these enhancers on different alleles would be consistent with a mosaic model for epigenetic regulatory marks in the population, rather than a poised state encoded by different marks within the same nucleosome.

Such a model also suggests that there is a relationship between DNA methylation and histone states at the candidate enhancers. One of our findings is that the mosaic candidate enhancers are enriched at what have been generally described as CpG island shores, the ±2 kb flanking CpG islands^31^. One such shore at the *GLT1* gene has previously been found to encode an enhancer that is, when silenced, enriched in DNA methylation and H3K27me3 but depleted in H4ac^40^. For this individual example it appears that the DNA methylation and histone modifications are associated with enhancer function, potentially representing enhancers more generally. The *GLT1* CpG island shore demonstrates co-localization of DNA methylation and H3K27me3, an association which has been shown to occur outside but not within CpG islands^41, 42^ using ChIP for H3K27me3 followed by bisulphite sequencing to detect the DNA methylation state on the alleles bearing the H3K27me3 nucleosomes. The more general finding that increased DNA methylation at CpG island shores correlates with decreased gene expression^31^ supports a model of these sites representing DNA methylation-sensitive enhancers where Polycomb-mediated H3K27me3 modification adds to the silencing of these regulatory elements.

We conclude that the epigenome of a cell population is, in fact, a collection of epigenomes, or a meta-epigenome, reflecting the presence of subpopulations of cell subtypes even in highly purified cell samples. This study’s use of CD34+ HSPCs was fortuitous, as this cell type is extremely well studied, and already recognized to be composed of multiple cell subtypes expressing the CD34 surface marker^33^. It is possible that the multipotent nature of the CD34+ HSPC cells is associated with greater epigenetic variability than more lineage-committed cell types, as suggested previously^43^. The prediction based on the observation of epigenetic variability in monocytes that HSPCs should be epigenetically variable between individuals^12^ is supported by our findings, although their findings could also be re-interpreted to be due to the long-term persistence of varying subtypes of monocytes in their studied populations. We propose that purified CD34+ HSPCs are not likely to be unique in being composed of multiple cell subtypes, and that detailed study of any presumed uniform cell population will reveal subpopulation heterogeneity. The results indicate that reference epigenomes based on the analysis of single or limited numbers of samples will contain epigenetically-variable loci at which marks are unstable, and the co-localisation of chromatin states at the same genomic position cannot reliably be taken to infer the presence of the epigenetic marks on the same alleles.

We show that the meta-epigenomic structure in the cell population can be exploited to estimate the number of cell subtypes present (using an approach like NMF^34, 35^) and their functional characteristics (by studying the properties of the genes located in proximity to the epigenetically variable *cis*-regulatory elements). Such meta-epigenomic analytical approaches could find an early application in cancer research. There are now consistent observations that epigenetic variability exists within cancer cells that have been extensively purified from contaminating cell types^44^ and even in non-neoplastic cervical epithelial cells in women who later develop cervical neoplasms^11^. These observations have been described to involve “stochastic variation” of epigenetic regulation^44^, but the CD34+ HSPC observations add a further layer of complexity, requiring that we understand cell subtype structure within the tested cell population before we can define any additional epigenetic variability as stochastic or disease-associated. This is a far-reaching issue, because while there now exist approaches that attempt to account for cell subtype heterogeneity within mixed cell populations tested using epigenome-wide assays, implicit in those approaches is that the subtypes of cells contributing to the variability can be distinguished histologically or by the use of cell sorting approaches^22, 45, 46, 47, 48^. What we show here is that even in histologically-identical, sorted cells, there exist subpopulation structures that continue to influence the results of epigenome-wide assays, and that the current approaches that rely on the use of sorted subpopulations of cells potentially do not go far enough to capture these influences.

The current study also has significant implications relevant to the interpretation of EWAS results. If a locus is found to change DNA methylation to the moderate extent typical of the results of non-cancer disease studies^5^, a distinction needs to be made between changes at loci that are normally invariant between individuals (those at the extremes of DNA methylation) and loci normally variable between individuals (intermediate methylation levels). In the former case, epigenetic changes must be occurring in some cells within the population studied. Changes at loci that are normally epigenetically variable may, however, be due to changes in cell subtype proportions between the individuals tested and not involve epigenetic changes occurring in any of the cells in the population tested. We increasingly recognize the influence of cell subtypes to be an issue in epigenomic studies of mixed cell types like peripheral blood leukocytes^22^ but the same concern arises even in purified cell populations, which are likely to have unrecognized cell subtype structure. The development of meta-epigenomics as a field of study is an essential early step towards improvement of our design and interpretation of the results of epigenome-wide assays.

## METHODS

The Supplementary Data section provides greater detail about the samples and methods used in this study. The sample collection component to the study was approved by our institutional review board (IRB) and was in accordance with Health Insurance Portability and Accountability Act (HIPAA) regulations. Cord blood samples were obtained at the time of delivery of healthy, non-anomalous neonates with normal growth based on birth weight and ponderal index nomograms. CD34+ HSPCs were purified from the cord blood using magnetic immunosorting, with purity confirmed by flow cytometry. DNA from the purified cells was used for DNA methylation assays, the HELP-tagging assay^24^ for genome-wide analysis and bisulphite PCR amplicon sequencing for verification and validation studies. HELP-tagging was performed on genomic DNA from the frozen CD34+ HSPCs, digested to completion by either HpaII or MspI, following which the digested DNA was ligated to two custom adapters containing Illumina adapter sequences, an EcoP15I recognition site and the T7 promoter sequence. Using EcoP15I, we isolated sequence tags flanking the sites digested by each enzyme, methylation-sensitive HpaII or methylation-insensitive MspI, followed by massively-parallel sequencing of the resulting libraries (Illumina technology). HpaII profiles were obtained for each sample (n=29), calculating methylation scores using a previously generated MspI human reference, which was also used to determine the degree of technical variability in the assay, using three replicates. For targeted bisulphite sequencing, we bisulphite-converted 200 ng of DNA using the Zymo EZ-96 Methylation-Lightning Kit. After separate PCR amplification of 10 target regions (primers listed in Supplementary Table 2), we pooled the amplicons in equal ratios and generated Illumina libraries using robotic automation (Tecan). In total, 15 libraries were multiplexed on the Illumina Miseq for 250 bp paired end sequencing. Bisulphite conversion efficiency was calculated as the percent conversion of cytosines in a non-CG context. Massively-parallel sequencing was performed using the Illumina HiSeq 2000 for HELP-tagging, and the Illumina MiSeq for bisulphite PCR amplicon studies.

Variability of DNA methylation was calculated using the Median Absolute Deviation (MAD) value, previously used to define variably-methylated regions^49^. The MAD calculation is less sensitive to outlying values, giving a more robust and conservative variance estimate. MAD values were calculated from our HELP-tagging data (29 neonates) and from reduced representation bisulphite sequencing (RRBS) data generated on the same cell type by the Roadmap in Epigenomics (7 adults).

We performed verification studies at 10 separate loci on 7 of the 29 samples tested using HELP-tagging, and for validation we added 8 new CD34+ HSPC samples from similarly healthy infants with normal growth. Using the DNA methylation value for each assayed HpaII locus, we calculated the MAD of DNA methylation for both validation and verification data sets, and observed an increase in variability when background technical variation (defined by control MspI HELP-tagging representations) was exceeded (Supplementary Fig. 2). The MAD distribution was calculated genome-wide, at HpaII sites overlapping common SNPs (minor allele frequencies ≥1%) and at the 10 bp immediately flanking these common SNPs (Supplementary Fig. 3). Kolmogorov–Smirnov testing was performed to measure the significance of differences of distributions of MAD values.

The youngest individual studied by the Roadmap in Epigenomics from whom RRBS, chromatin and transcriptional studies had been performed was chosen for further analysis. Wiggle track data representing DNase hypersensitivity and ChIP-seq for H3K4me3, H3K4me1, H3K9me3, H3K27me3, H3K27ac and H3K36me3 were collected from the Roadmap in Epigenomics web resource (http://www.roadmapepigenomics.org/data). All wiggle tracks were converted to bigwig format using the UCSC Genome Browser utility *wigToBigWig* version 4 (http://hgdownload.cse.ucsc.edu/downloads.html). Subsequently, the utility available through the UCSC Genome Browser *bigWigAverageOverBed* (http://hgdownload.cse.ucsc.edu/downloads.html) was used to calculate the sum of the ChIP-seq signals over 100 bp genomic intervals spanning the 22 autosomal and 2 sex chromosomes, a resolution which we believe is sufficient to characterize chromatin states, being smaller than an individual nucleosome while usually including no more than 0–1 HpaII sites per window. ChIP-seq signals summed over 30,956,785 intervals were generated and formatted in bedGraph format.

The Roadmap in Epigenomics data are provided as raw signals and not as defined peaks. To avoid imposing excessive processing on these data, we used as simple and intuitive an approach as possible. The ChIP-seq bedGraph files were log-transformed to exaggerate highly-positively skewed signal density, a prerequisite for the recursive kernel density learning framework for robust foreground object segmentation approach^29^. This image processing technique relies on removing background until the remaining signal is bimodal and approximately Poisson distributed. Gaussian kernel density was estimated using the *density* function in R resulting in multiple modes of signal density that are increasingly smaller. Because signal intensity was derived from a ChIP-seq read count, the measure should be an approximate Poisson distribution, and we aimed to eliminate low signal intensity signal modes iteratively. Using the *turnpoints* function within the *pastecs* library in R, we recursively identified the modes of signal intensity and set signal thresholds based on the maximum mode, which was also generally the leftmost mode. The algorithm ran until at most 2 signal modes remained and the resulting distributions were approximately Poisson distributed. The results are shown in **Figure S4**, with the stepwise approach for the H3K4me1 signal illustrated as an example of the process.

100 bp windows were then classified as having the chromatin state or not. Windows of chromosome 1 were used to train the Segway algorithm^28^, which then annotated 7 features genome-wide. A subset of 100 bp windows containing HpaII sites tested using HELP-tagging was chosen for self-organising map (SOM) construction^30^. All SOM analysis was completed using the Java SOMToolbox from Institute of Software Technology and Interactive Systems at the Vienna University of Technology (http://www.ifs.tuwien.ac.at/dm/somtoolbox/). Out of 30,956,785 100 bp genomic intervals, 1,520,684 intervals overlapping 1,696,696 HpaII sites were chosen to reduce the dimensionality of the dataset and greatly reduce the required computation. An input data matrix was created, where rows represent the 1,520,684 vectors defined by genomic intervals and the columns represent the observations for the investigated tracks (*i.e.* processed ChIP-seq signal). In total, two SOMs were constructed, one representing the chromatin states from the Roadmap in Epigenomics, the other the ChromHMM annotation^32^, choosing a map size to yield 100–200 interval-vectors per map unit to reduce the required computation while generating maps of sufficient resolution to aid in further analysis. For all SOMs the standard SOM algorithm by Kohonen was employed, using the default SOMToolbox settings of learnrate=0.7 and randomSeed=11. Java code was implemented over approximately 120 hours using a high performance computing (HPC) cluster and 100 GB of virtual memory.

We performed an analysis to define CD34+ HSPC genomic annotations using the Segway genomic segmentation approach^28^. Using the *segway* package, annotations were generated from the 7 processed chromatin state signal bedGraph files. A Segway segmentation of the genome was created by training on chromosome 1, using the results to annotate the whole genome by requesting 7 labels, allowing each chromatin state signal to vary independently from the others. Furthermore, we required at least 1,000 bp segments, a 500 bp ruler and a 500 bp ruler scale, which had the effect of smoothing across the segmentation, which we found to be excessively sensitive to varied signal at the default settings. Segway was completed requesting 10 simultaneous runs, over which maximum likelihood estimations regarding chromatin state were performed. Segway code was implemented using the HPC cluster with the training step taking approximately 72 hours and the identify step taking approximately 2 hours using 40GB of virtual memory.

To interpret our genomic annotations (Segway features) and epigenomic variability (DNA methylation MAD values), we created contour plots plotting enrichment within map units within the SOMs. The enrichment within SOMs was determined by a proportion test, specifically asking whether the observed proportion of a feature within a map unit was significantly greater than the expected proportion given the distribution of Segway features overall. A cutoff of 23.09, the 98.5^th^ centile of overall MAD, was used in order to dichotomize MAD into high and low variable states (for reference, note that ln(23.09) = 3.14, the cutoff shown in Supplementary Fig. 2). From calculated proportions of highly variable 100 bp intervals within the SOM units, we performed a proportion test, specifically testing whether the observed proportion of a highly variable interval within a map unit was significantly (alpha=0.05) greater than the expected proportion, given the overall proportion of highly variable intervals. The *MASS* library in R along with the *kde2d* and *contour* functions were used to represent the density of variability-enriched map units with a contour plot over SOMs.

We analyzed as metaplots RefSeq gene and CpG island annotations and the 10 kb regions flanking these annotations, allowing us to study the relationship between MAD values and these genomic elements. The bodies of RefSeq genes and CpG islands were split into deciles in order to be able to compare genes of varying lengths, and the 10 kb flanking region was separated into 100 bp windows. Gene coordinates were rounded to the nearest 100 bp to ensure that sequences were not represented twice. Segway features were divided into 100 bp intervals and matched with 100 bp windows or matched with RefSeq gene deciles, allowing more than one Segway feature to match a particular window or decile. For each 100 bp window or decile, the frequency of each Segway feature was calculated and plotted.

To test the relationship of the Segway-derived functional annotations and DNA methylation variability with transcription, we used the CD34+ HSPC RNA-seq data from the same individual from whom chromatin data were obtained (RO_01549, Supplementary Table 4, GEO accession number SRA010036). Of the 17 available RNA-seq runs, we used SRR453391, corresponding to 16,000,000 reads and 2.4 gigabasepairs of sequence. Reads were quality controlled using *FASTX-Toolkit* v0.0.13, with *fastq_quality_trimmer* trimming nucleotides with quality lower than 3 and removing sequences shorter than 17 bp. Reads were aligned to the human genome using *GSNAP* version 2012-07-20 requesting at most 10 alignments for multiple aligned reads. *SAMtools* v0.1.8 was used to convert the alignment to BAM format. The *Cufflinks* v2.02 program *Cuffdiff* v2.0.2 was used to calculate fragments per kilobase per million reads (FPKM) values for RefSeq genes, employing normalization by the upper quartile of the number of fragments mapping to individual loci and default weighting of multiply aligned reads based on the number of alignments. FPKM values were used to separate genes into those that were not expressed (8,963 genes) and those that were expressed by quartile of expression (7,872 genes per quartile). Expression information was then linked to the annotated RefSeq gene body deciles and 10 kb flanking regions, thus allowing the stratification of Segway features overlapping 100 bp windows and gene body deciles by gene expression.

DNA methylation variability (MAD values), Segway features and gene expression levels were studied relative to RefSeq genes, dividing the bodies of the genes into deciles to allow comparisons for genes of different sizes, and extending the analysis using the 100 bp windows to flank the gene body 10 kb upstream and downstream. A similar approach to study the margins and flanking regions of CpG islands was also performed.

We tested whether the peak loci for enrichment of 100 bp windows for features 4 and 6 reached statistical significance. The peak window for feature 4 is at −1,500 bp upstream from RefSeq transcription start sites, where it comprises 31.50% of the features (compared with 15.28% genome-wide), while feature 6 is at peak enrichment at −100 bp, comprising 56.60% of annotated features (compared with 9.27% genome-wide). A one way proportion test for each feature at the peak locations shows significance for enrichment for both features (p<0.001). We then tested whether the variability of DNA methylation at these windows of peak feature enrichment was also significantly increased. We compared the MAD values for each feature at these peak enrichment sites with those values at the same number of windows randomly selected from either RefSeq genes, showing with a one way Wilcoxon rank sum test that variability for DNA methylation at these loci was also significantly increased for features 4 and 6 (p<0.001). Using the same analytical approach, feature 6 was found to be significantly enriched within CpG islands and feature 4 in the ±2 kb CpG island shores (p<0.001). We also tested whether the observed trend of increased DNA methylation variability at feature 4 associated with decreased gene expression levels was significant using the non-parametric Jonckheere trend test. The trend was significant at p=10^−8^.

To interpret DNA methylation variability observations, we asked the question whether the increased variability we observed at candidate promoters and enhancer sequences (Segway features 6 and 4 respectively) was occurring non-randomly at genes with known functions. For each RefSeq gene, Segway feature 6 (promoters) overlapping the TSS and Segway feature 4 (enhancers) occurring within 5 kb flanking the TSS were isolated. We calculated the median MAD over these features. The MAD over promoters and enhancers was dichotomized using the 23.09 value (98.5^th^ percentile) allowing genes to be characterized as having high and low variability over both promoters and enhancers. The Broad Institute’s Gene Set Enrichment Analysis web applet (http://www.broadinstitute.org/gsea/) performs a hypergeometric/Fisher’s exact test on gene list supplied from the Molecular Signatures Database v3.1 (http://www.broadinstitute.org/gsea/msigdb/), to identify pathways differentially enriched for high and low variability enhancers using an False Discovery Rate (FDR) q value of less than 0.05 (Supplementary Table 6). We further demonstrate this association using Reactome pathways isolated through the Pathway Commons (http://www.reactome.org/static_wordpress/about/ and http://www.pathwaycommons.org). Gene pathways were visualized in Cytoscape v2.8.3 with edges representing the physical interactions between nodes stored in the GeneMANIA v3.2 plugin. DNA methylation variability was high for promoters for both housekeeping (KEGG Ribosome) and leukocyte-specific (Leukocyte Transendothelial Migration) genes at candidate promoters (Segway feature 6), but substantially decreased or absent at housekeeping and not leukocyte-specific genes at candidate enhancer loci (Segway feature 4, Supplementary Figure 11).

To infer the number of cell subtypes present in the CD34+ HSPC population, we used the non-negative matrix factorization (NMF) approach that has previously been applied to transcriptomic data^34, 35^. NMF has been employed successfully in deconvolving gene expression data^35^, but it has not previously been applied to DNA methylation datasets. The goal of an NMF algorithm is to deconvolve a matrix *V*, a (n × p) matrix, in order to find an approximation the matrices *W* and *H* such that:

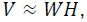

where W and H are (n × r) and (r × p) non-negative matrices. The rank of the matrix (r) should be greater than 0 and at least 2 to represent 2 subpopulations. Because the methylation outcome is binary, and the existing NMF algorithms allow DNA methylation to vary without this constraint the estimated matrix, W′ may not be interpretable directly, and improvement of the technique may allow the methylation pattern of the individual subpopulations to be estimated. We applied an existing NMF algorithm to understand the presence of subpopulations within our dataset but did not interpret the specific values within W′, rather we focused on the difference between the actual and simulated datasets.

Utilizing the R package *deconf*, we varied the matrix rank and estimated the matrices W′ and H′ such that:

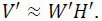

The distance between the original dataset V and V′ was calculated as the Frobenius norm:

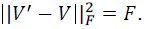

The process was repeated 100 times per value of r, subsetting the data to 10,000 HpaII sites in order to make the algorithm computationally tractable. A plot of the distribution of *F* as a function of increasing r (cell subpopulations) is shown (Figure 6). In addition, the distribution of Frobenius norms for each cell subpopulation was compared to the preceding cell subpopulation with a two sample t-test, testing for a difference in distribution of Frobenius norms between successive simulation levels. Because estimating additional subpopulations will always explain additional variability, a smooth spline was fit to the data to look for inflection points indicating a local minimum in the Frobenius norm when related to cell subpopulations.

Further detailed descriptions of all of the analyses performed are provided in the Supplementary Data.

## END NOTES

### Accession codes

All HELP-tagging data generated are deposited into the Gene Expression Omnibus (GEO) database under accession number GSE49836.

## Acknowledgements

Support for this project was provided by Roadmap Epigenomics R01 HD063791 (Einstein/Greally). Support was also provided by Einstein’s Medical Student Training Program award (to NAW, NIH NIGMS T32 GM007288), and Einstein’s Center for Epigenomics, including the Epigenomics Shared Facility and Computational Epigenomics Group.

## Author contributions

NAW, FD: performed experiments, analyzed data, wrote manuscript. YMZ: performed experiments. JCM, AG: provided guidance and performed analytical approaches. FHE, JMG: designed study, analyzed data, wrote manuscript.

## Additional information

Competing financial interests:

The authors declare no competing financial interests.

